# Diffusion-weighted magnetic resonance spectroscopy enables cell-specific monitoring of astrocyte reactivity *in vivo*

**DOI:** 10.1101/350306

**Authors:** Clémence Ligneul, Edwin Hernández-Garzón, Marco Palombo, María-Angeles Carrillo-de Sauvage, Julien Flament, Philippe Hantraye, Emmanuel Brouillet, Gilles Bonvento, Carole Escartin, Julien Valette

## Abstract

The diffusion of brain intracellular metabolites, as measured using diffusion-weighted magnetic resonance spectroscopy *in vivo*, is thought to specifically depend on the cellular structure constraining them. However, it has never been established that variations of metabolite diffusion, e.g. as observed in some diseases, could indeed be linked to alterations of cellular morphology. Here we demonstrate, in a mouse model of reactive astrocytes, that advanced diffusion-weighted magnetic resonance spectroscopy acquisition and modeling techniques enable non-invasive detection of reactive astrocyte hypertrophy (increased soma radius, increased fiber radius and length), as inferred from variations of myo-inositol diffusion, and as confirmed by confocal microscopy *ex vivo*. This establishes that specific alterations of intracellular metabolite diffusion can be measured and related to cell-specific morphological alterations. Furthermore, as reactive astrocytes are a hallmark of many brain pathologies, this work opens exciting perspectives for neuroscience and clinical research.

## Main text

The brain is an organ of immense complexity, whose function relies on the interactions of different cell types such as neurons and astrocytes, each with their own morphological and metabolic specificities. The quest for non-invasive tools capable of probing cellular status to better understand the normal brain, but also to characterize and monitor its dysregulations under pathological conditions, has been driving considerable methodological research. In particular, over the last 30 years, nuclear magnetic resonance has delivered revolutionary tools to non-invasively probe brain cell structure and metabolism. Magnetic resonance imaging (MRI), which relies on the detection of water molecules, is now widely used to investigate brain characteristics (anatomy, function, microstructure and metabolism) *via* several types of contrasts. This includes diffusion MRI, which exploits the fact that water diffusion is sensitive to restrictions imposed by cellular membranes, thus indirectly informing on the underlying cellular structure. In parallel, magnetic resonance spectroscopy (MRS), where the signal originates from metabolites resonating at frequencies other than water’s, offers unparalleled access to cellular metabolism. MRI benefits from good sensitivity due to the abundance of water molecules, thus allowing high spatio-temporal resolution, but suffers from its poor cellular specificity, since water molecules are found in all cell types as well as in the extracellular space. This is particularly problematic for diffusion MRI, which consequently cannot disentangle the contribution of the different cell types (even though it is often assumed, more or less explicitly, that neuronal structure is the main determinant of water diffusion, since neurons represent the largest cellular volume fraction in the brain (Hertz, 2008)). This contrasts with MRS which, despite its low sensitivity due to low metabolite concentrations, benefits from exquisite specificity. Indeed, most brain metabolites are confined in the intracellular space, and some of them are even considered to be cell-specific: N-acetylaspartate (NAA) and glutamate (Glu) have for example been reported to be mostly found in neurons, while choline compounds (tCho) and myo-inositol (Ins) are thought to be predominantly in glial cells (Choi et al., 2007; Gill et al., 1989) (more particularly in astrocytes for myo-inositol (Brand et al., 1993)). Hence, the diffusion of these metabolites, as measured using diffusion-weighted MRS (DW-MRS), must specifically depend on the cellular structure constraining them (Palombo et al., 2017c). Some recent works, including ours, support this idea: indeed, different diffusion behaviors that can be reasonably (and quantitatively) explained in terms of different cellular structure have been reported for neuronal and glial metabolites in the healthy brain (Palombo et al., 2017c). However it has never been established that variations of metabolite diffusion, e.g. as observed in some diseases (Ercan et al., 2016; Wood et al., 2012), could indeed be linked to alterations of cellular morphology.

The present work aims at demonstrating, for the first time, that DW-MRS allows probing cell-specific structural alterations. To achieve this goal, we chose to study a mouse model of astrocyte reactivity induced by the cytokine ciliary neurotrophic factor (CNTF). For a long time described as supporting actors in normal brain function, with basic roles in physical and metabolic scaffolding of neurons, astrocytes are emerging as key players in transmission through neuronal networks, *via* fine modulation of synapsis, synaptic connectivity establishment and spread of synaptic signalization (Araque et al., 2014; Navarrete and Araque, 2014). Moreover, given that almost all brain functions are essentially regulated by glial cells, there is an increasing awareness of their importance in all spectrum of neurological disorders (Zuchero and Barres, 2015). Glial cells are recognized as first responders against external stimuli, and the first cells affected when neurodegenerative diseases or cerebrovascular accidents appears. Reactive astrocytes are a hallmark of many brain pathologies (Dossi et al., 2018), including neurodegenerative diseases, and have been proposed to play a major role in synaptic plasticity (Liddelow and Barres, 2017; Tyzack et al., 2014). They are characterized by the overexpression of the glial fibrillary acidic protein (GFAP) and vimentin, increased release of cytokines and chemokines, and hypertrophy. In the context of pure astrocyte reactivity as induced in the model we propose to investigate, with neither neuronal death nor microglial activation (Escartin et al., 2006; Escartin et al., 2007), our hypothesis is that astrocytic metabolites would exhibit diffusion alterations that are quantitatively consistent with astrocytic hypertrophy as measured by confocal microscopy, while other metabolites, in particular neuronal metabolites, would exhibit negligible or no diffusion alteration. This would establish that specific alterations of intracellular metabolite diffusion can be measured and related to cell-specific morphological alterations. As a more immediate application, this would open fantastic possibilities to monitor astrocytes in various contexts, from plasticity to disease.

## RESULTS

### MRS identifies neuronal and glial metabolic modifications associated with reactive astrocytes

Astrocyte reactivity was induced in mice by intra-striatal injection of lentiviral vectors specifically targeting neurons and encoding for the human ciliary neurotrophic factor (CNTF), or β-galactosidase (LacZ) for control (**Fig. 1A**). All DW-MRS measurements were performed at 11.7 T with a cryoprobe, using an optimized stimulated-echo sequence(Ligneul et al., 2017). We measured the metabolic profile of both phenotypes from spectra acquired at the lowest diffusion-weighting and shortest diffusion time. For visual inspection, the sum of spectra acquired in each individual mouse for both groups is shown in **Fig. 1B**, showing stable absolute levels of total creatine, but variations of concentration for many other metabolites. In the CNTF group, we report a lower concentration of NAA (−20%), glutamate (−17%) and taurine (−18%), and a strong increase in myo-inositol concentration (+92%) (**Supplementary Table S1**). Lactate concentration may seem to slightly increase, but without significance. This metabolic remodeling is consistent with a previous MRS study conducted on rats injected with lenti-CNTF (Carrillo-de Sauvage et al., 2015), except for choline compounds that were increased in rats while they remain stable here. The macromolecule baseline is very similar between CNTF and control mice (**Fig. 1C**), except for an additional resonance around 1.2 ppm, likely corresponding to β-hydroxybutyrate. It is worth noting that the amplitude of macromolecule signal perfectly matches between both groups (except for this additional resonance at 1.2 ppm), underlining the absence of massive cellular loss or genesis in the MRS volume of interest, in line with the stability of striatal volume as measured from T2-weighed MRI images (31.6±2.0 mm^3^ in CNTF group *versus* 31.4±1.6 mm^3^ in control group).

**Figure 1.**
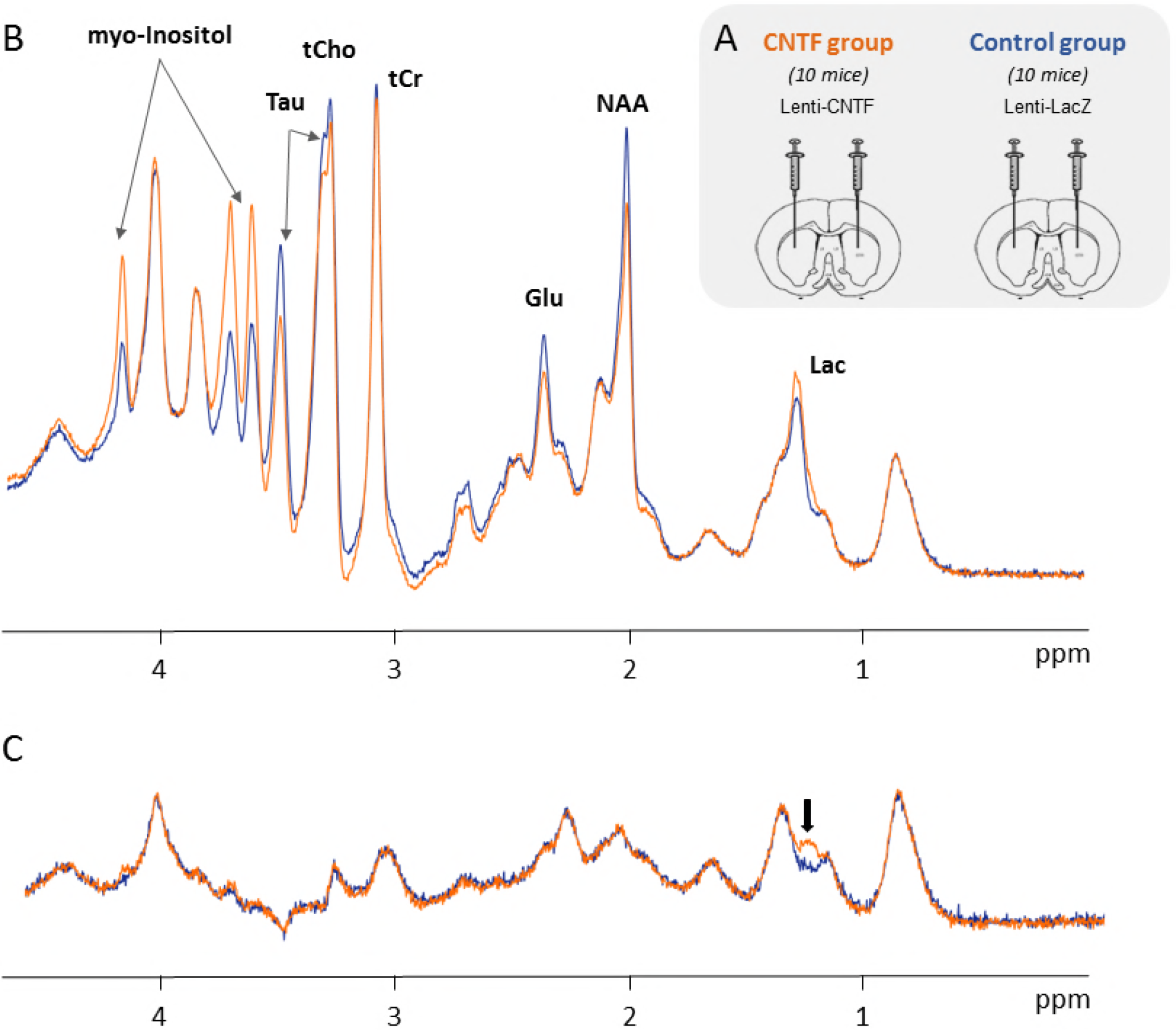
Alterations detected in the CNTF mouse model of reactive astrocytes using *in vivo* MRS. (**A**) Astrocyte reactivity was induced in the striatum by bilateral injection of lentiviral vectors specifically targeting neurons and encoding for CNTF, or β-galactosidase (LacZ) for control. (**B**) Average of all spectra acquired in each group (10 mice per group) to illustrate variations of metabolite levels (Glu=glutamate, Lac=lactate, Tau=taurine, tCho=choline compounds, tCr=total creatine). (**C**) Average of macromolecule spectra acquired in each groups. These spectra are very similar except for the small peak appearing at ~1.2 ppm, likely corresponding to β-hydroxybutyrate. Neither scaling nor filtering was applied to spectra.

### DW-MRS reveals myo-inositol as a specific diffusion marker of astrocyte reactivity

In recent works performed in healthy animals, we have proposed that high diffusion-weighting DW-MRS data primarily reflect the radius of brain cell fibers (dendrites, axons, astrocytic processes…) (Palombo et al., 2017b), while long diffusion times data primarily reflect fiber long-range structure, in particular complexity and length (Palombo et al., 2016). Here we measured signal attenuation of reliable metabolites (CRLB<5 %) up to a diffusion-weighting *b*=50 ms/μm^2^, at fixed diffusion time *t*_*d*_=53.2 ms, which is well suited to probe restriction in fibers’ transverse plane. Among the six intracellular metabolites measured reliably, only myoinositol exhibited significantly different signal attenuation curves from control (**Fig. 2B**), with stronger attenuation in the CNTF group, suggesting that myo-inositol is diffusing inside fibers of larger radius. One can notice slightly stronger signal attenuation for tCho and slightly weaker NAA signal attenuation in the CNTF group, but these trends are not significant. As can be seen on **Fig. 2A**, macromolecules contribute a lot to high-*b* spectra: the intensity of the broad macromolecules peak at 0.9 ppm is comparable to NAA intensity. Therefore, it is crucial to properly quantify their signal for such experiments (Palombo et al., 2017c). Here the excellent spectral quality and signal-to-noise ratio SNR at all diffusion-weightings (at *b*=50 ms/μm^2^ the SNR, calculated relative to NAA, is 26 ± 2 in the CNTF group, and 30 ± 2 in the control group), as well as for macromolecules (**Fig. 1C**), allow reliable evaluation of the signal attenuation.

**Figure 2.**
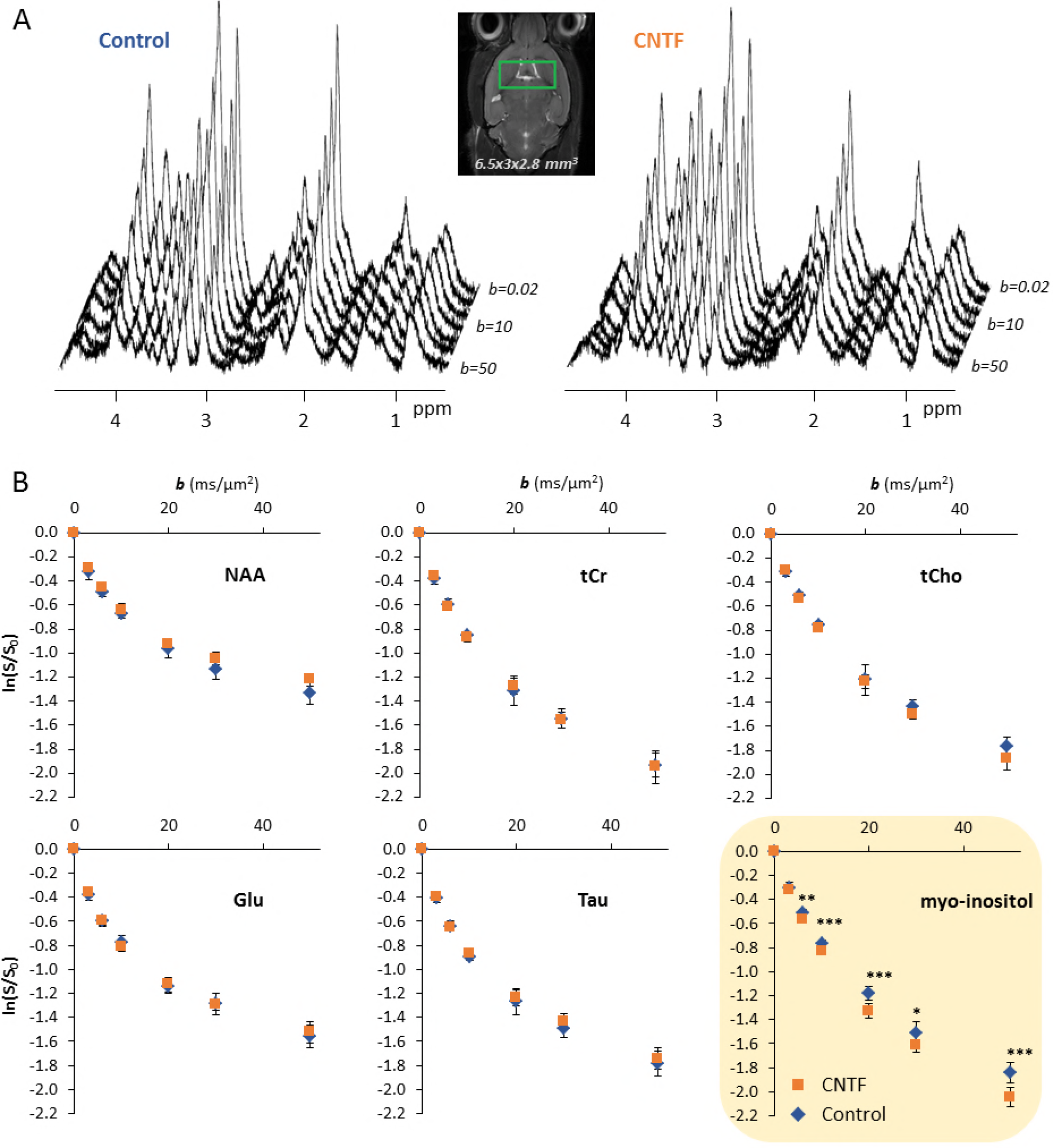
DW-MRS data at high diffusion-weightings in the CNTF mouse model of reactive astrocytes. (**A**) Spectra acquired up to high diffusion-weighting (from *b*=0.02 ms/μm^2^ to 50 ms/μm^2^, at fixed *t*_*d*_=53.2 ms) in one control mouse (left) and one CNTF mouse (right). (**B**) Signal attenuation as a function of diffusion-weighting *b* for all six intracellular metabolites reliably quantified, in both groups (blue diamond=control group; orange squares=CNTF group). Data points and error bars stand for mean±s.d. (*n*=10 mice per group). Only myo-inositol exhibits significantly different diffusion behavior (ANOVA group\**b* interaction: p=0.0048). Unpaired Student’s t-tests were run as *post hoc* tests for each *b* to assess significantly different data points between groups (*: p<0.05, **: p<0.01, ***: p<0.001, after Bonferroni correction). (Glu=glutamate, Tau=taurine, tCho=choline compounds, tCr=total creatine).

We then measured metabolite apparent diffusion coefficient (ADC) at long *t*_*d*_ (up to *t*_*d*_=2003.2 ms), leaving time for metabolites to explore the long-range fiber structure (**Fig. 3**). Again, only myo-inositol exhibited significant variation, with ADC decreasing less in the CNTF group as *t*_*d*_ is increased (**Fig. 3B**), indicating less long-range obstacles (which at first glance can be qualitatively interpreted as “longer fibers”). Spectral quality and SNR remain acceptable even in the least favorable situation (SNR=28 ± 4 in CNTF group and 30 ± 4 in control group at *t*_*d*_=2003.2 ms and *b*=3.6 ms/μm^2^, calculated relative to NAA).

**Figure 3.**
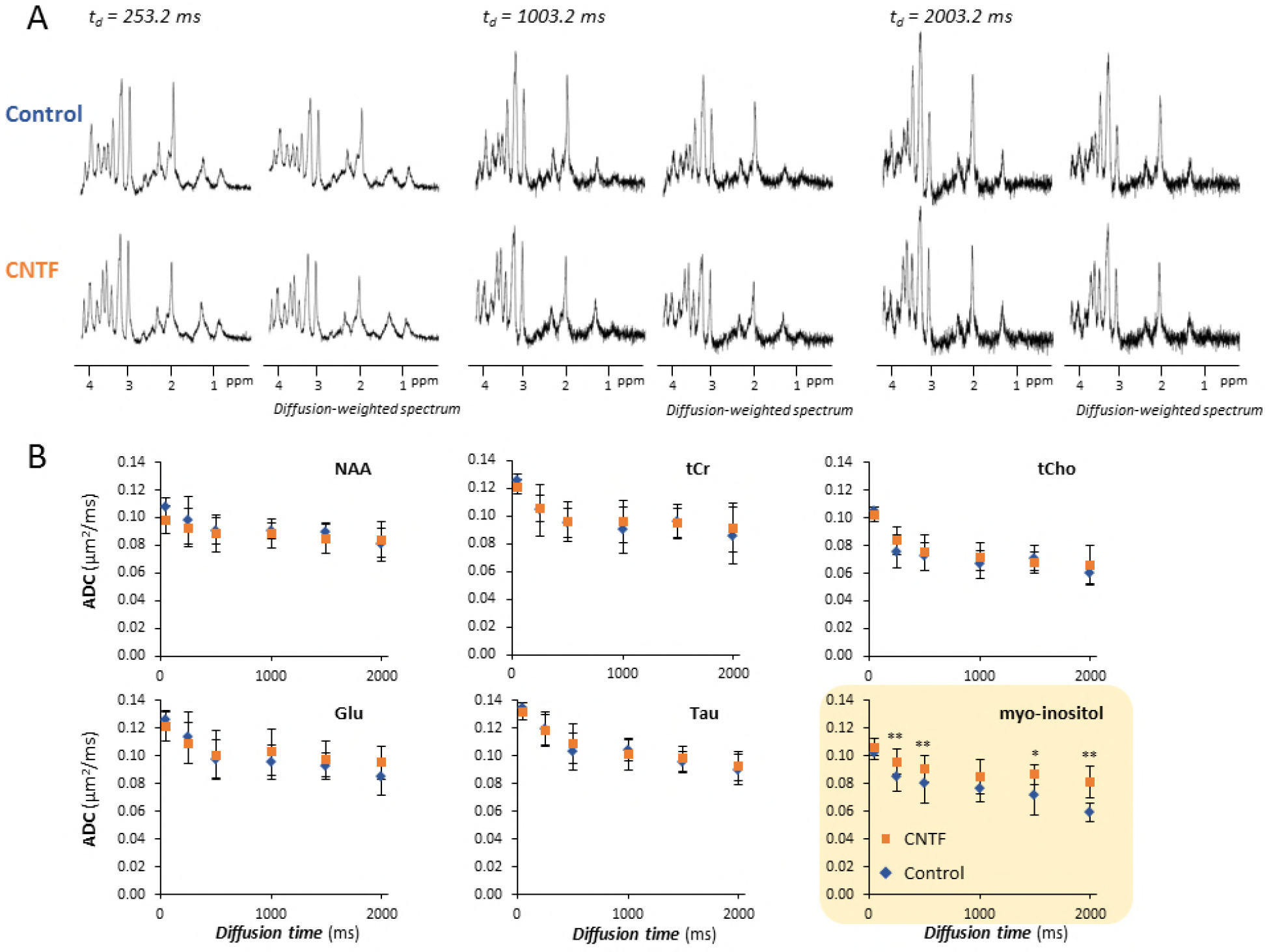
DW-MRS data at long diffusion times in the CNTF mouse model of reactive astrocytes. (**A**) Spectra acquired in a control mouse and a CNTF mouse at long diffusion times, without and with diffusion-weighting. Note that the exact *b*-values are given in **SI Appendix**, **Table S5**. (**B**) Apparent diffusion coefficient (ADC) as a function of the diffusion time (*t*_*d*_) for all six intracellular metabolites reliably quantified, in both groups (blue diamond=control group; orange squares=CNTF group). Data points and error bars stand for mean±s.d. (*n*=10 mice per group). Only myo-inositol exhibits significantly different diffusion behavior (ANOVA group\**b* interaction: p=0.016). Unpaired Student’s t-tests were run as *post hoc* tests for each *t*_*d*_ to assess significantly different data points between groups (*: p<0.05, **: p<0.01, after Bonferroni correction). (Glu=glutamate, Tau=taurine, tCho=choline compounds, tCr=total creatine).

In the end, both high-*b* and long-*t*_*d*_ experiments reveal specific variations of myo-inositol diffusion in the CNTF group, which are qualitatively consistent with diffusion inside thicker and longer fibers, as *a priori* expected for hypertrophic reactive astrocytes. Note that we also performed DW-MRI measurements, but no significant variation of related metrics (mean diffusivity, fractional anisotropy and mean kurtosis) were detected in the striatum (**Supplementary Fig. S1**).

### Diffusion modeling allows quantifying structural alterations of myo-inositol diffusion compartment

We then attempted to model myo-inositol diffusion data to quantitatively evaluate structural alterations of myo-inositol compartment (**Fig. 4**). High diffusion-weighting data were analyzed using analytical models of diffusion in randomly oriented cylinders (Palombo et al., 2017b), slightly modified to also account for diffusion in spherical soma representing 20% of the total cell volume (Palombo et al., 2018a, b). We extracted fiber radii of 0.58±0.02 μm for the control group *versus* 0.85±0.02 μm for the CNTF group (+47%), and soma radii of 3.80±0.06 μm for the control group *versus* 4.66±0.07 μm for the CNTF group (+23%) (Supplementary Table S2).

**Figure 4.**
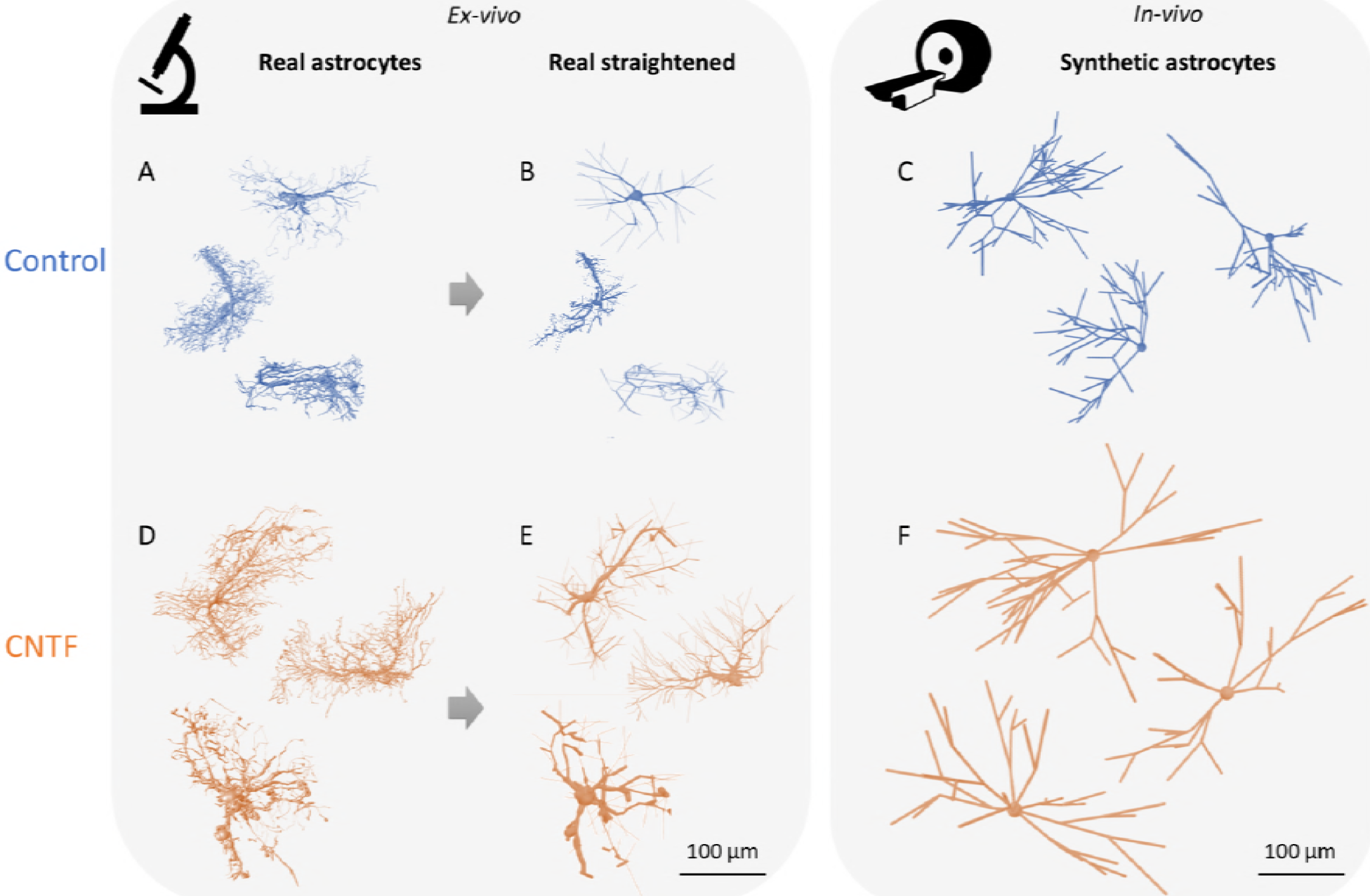
Comparison of some astrocytes obtained from direct *ex vivo* measurement by confocal laser scanning microscopy (“real” astrocytes) in control (**A**) and CNTF (**D**) mice; and from modeling of *in vivo* myo-inositol DW-MRS data (“synthetic” astrocytes) in control (**C**) and CNTF (**F**) mice. (**B**) and (**E**) represent typical astrocytes reconstructed from confocal microscopy after simplification of their structure (“real straightened” astrocytes, see Methods), from which we measure morphological parameters for quantitative comparison with synthetic astrocytes. In both real straightened and synthetic astrocytes, increased soma radius, fiber radius and fiber length are measured in CNTF (see main text and **SI Appendix**, **Tables S3** and **S4** for parameter values). To generate the representations for synthetic astrocytes in (**C**) and (**F**), some cell-graphs as extracted from long-*t*_*d*_ data were “thickened” by conferring some non-zero radius to their fibers and soma, corresponding to the best radii extracted from high-*b* data for each group.

Long *t*_*d*_ myo-inositol data were analyzed using a machine learning approach, elaborating upon the framework of numerical simulations in branched structures as we introduced recently (Palombo et al., 2016). Briefly, cellular fibers are modeled as mono-dimensional branched objects embedded in a three-dimensional space, dubbed “cell-graphs”. Here only two morphometric parameters were let as free parameters: the average number of successive embranchments (bifurcations) N_branch_ along each process, and the average segment length L_segment_ for a given segment of process comprised between two successive branching points. For each of these statistics, a Gaussian distribution was assumed. To generate the dictionary, a large set of different synthetic cells was generated for any given set of morphometric statistics values, and diffusion of many particles was then simulated in each cell according to a Monte Carlo algorithm. The corresponding diffusion-weighted signal was computed using the phase accumulation approach and summed over the whole set of cell-graphs to obtain the coarsegrained ADC. Random forest regression was then performed to compare experimental ADC time-dependency to this dictionary and thus estimate N_branch_ and L_segment_ for myo-inositol compartment. At this point we should mention that conventional cell analysis tools do not provide a direct estimation of N_branch_, which is an *ad hoc* parameter used to control the branching complexity of our cell-graphs. To facilitate further comparison with real astrocytes as measured by microscopy (see next section), we found it more convenient to generate many cell-graphs corresponding to the set of (N_branch_, L_segment_) values best explaining the data, and then, as for real cells, analyze these cell-graphs using the TREES Matlab toolbox to extract the “Branch Order” parameter characterizing the branching complexity (Cuntz et al., 2010). In the end, data modeling yields a preserved Branch Order in CNTF, but increased length of fiber segments between each embranchment, from 29±3 μm to 59±6 μm (+103%), resulting in overall increased fiber length (**Fig. 4** & **Supplementary Table S3**).

### *Ex vivo* microscopy confirms astrocytic hypertrophy in line with alterations of myo-inositol diffusion

We then confirmed astrocytic hypertrophy using confocal laser scanning microscopy (**Fig. 4** and **Supplementary Fig. S2** to **S5**). On reconstructed GFP-expressing astrocytes, we measured fiber and soma radii of respectively 0.50±0.08 μm and 3.7±1.2 μm for the control group *versus* 0.60±0.13 μm and 4.9±0.9 μm for the CNTF group (+20% and +32%). These data confirm thicker astrocytic processes for reactive astrocytes, and are in very good quantitative agreement with values extracted from modeling high-*b* myo-inositol data.

To analyze the long range-morphology of GFP-expressing astrocytes reconstructed by confocal microscopy, one should keep in mind that we are seeking a relevant comparison with synthetic cell-graphs estimated from myo-inositol long-*t*_*d*_ data (the “synthetic astrocytes”). Hence it is necessary to work with objects exhibiting commensurate levels of details. To do so, prior to the morphology analysis with the TREES toolbox, cells reconstructed from confocal microscopy images were simplified, essentially by removing small secondary structures that are “invisible” in long-*t*_*d*_ DW-MRS as they don’t induce ADC time-dependency at long *t*_*d*_ (Palombo et al., 2017a), and by “straigthening” undulating segments. In the end, after that simplification, we found that branching complexity did not change significantly, but we measured increased segment length between embranchments, from 15±5 μm to 25±7 μm (+67%). These data confirm longer astrocytic processes for reactive astrocytes, in agreement with the increased length estimated from modeling long-*t*_*d*_ myo-inositol data (**Supplementary Fig. S6** and **Supplementary Table S3**). It appears that estimates of L_segment_ in synthetic cells are about two times larger than real ones. However, despite that discrepancy in terms of absolute length, the overall agreement between synthetic and real astrocytes remains striking: L_segment_ statistically increase in CNTF compared to LacZ, while Branch Order tends to decrease, even if not significantly.

### Lactate diffusion as a probe of lactate cellular compartmentation?

So far we have reported the diffusion of intracellular metabolites and interpreted it in terms of cell structure; however, CNTF-induced astrocyte reactivity is not only associated with morphological changes, but also with energy metabolism alterations. In particular, increased oxidative metabolism has been reported in astrocytes (Escartin et al., 2007). Very intriguingly, we observed massive variation of lactate diffusion, which are visually apparent event just looking at spectra (**Supplementary Fig. S7**), lactate attenuation as a function of *b* being much weaker in the CNTF group (**Fig. 5A**). Note that lactate signal was too low for reliable determination at long *t*_*d*_, so that only lactate signal attenuation at *t*_*d*_=53.2 ms is reported here.

**Figure 5.**
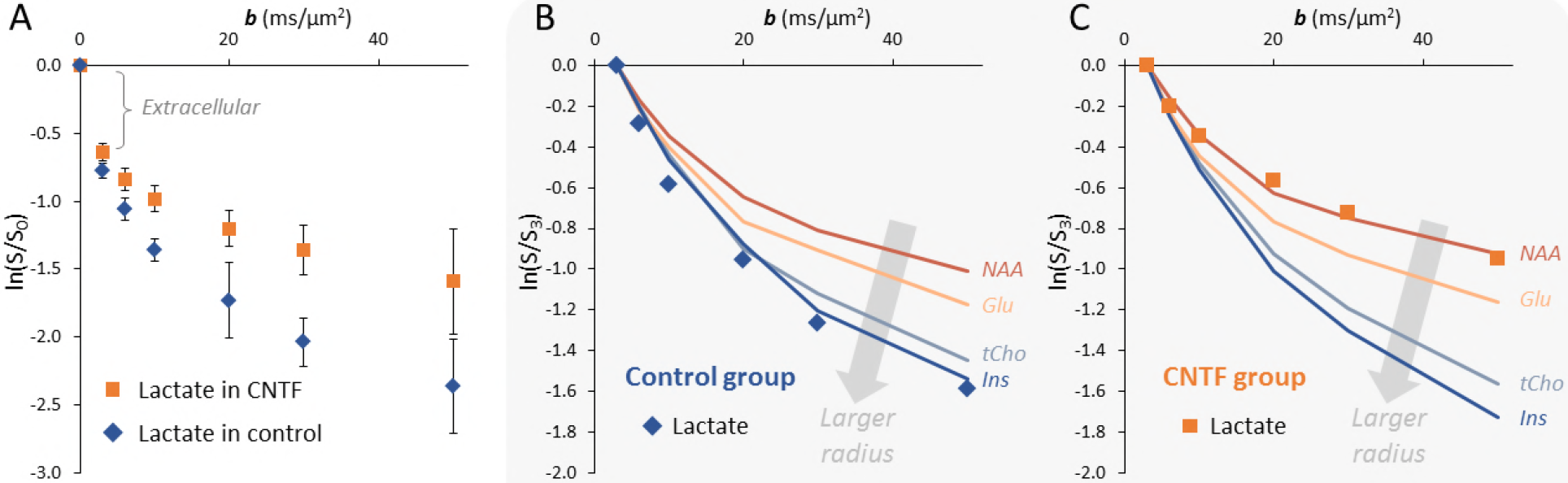
Massive variation of lactate diffusion suggests shift from predominantly astrocytic to predominantly neuronal compartmentation of lactate. (**A**) Lactate attenuation at high diffusion-weighting is much weaker in CNTF, indicating more restriction to diffusion. Error bars stand for s.d. (*n*=10 mice per group). (**B**) In control, when normalizing signal relative to *b*=3 ms/μm^2^ to minimize extracellular lactate contribution, lactate exhibits strong signal attenuation similar to myo-inositol, corresponding to diffusion in fibers of large radius (typical of diffusion within astrocytes). (**C**) In contrast, in CNTF, lactate diffusion is similar to NAA, corresponding to diffusion in fibers of small radius (typical of diffusion within neurons). (Glu=glutamate, Ins=myo-inositol, tCho=choline compounds).

Interpreting lactate diffusion is difficult because, unlike other metabolites, lactate has a significant extracellular fraction. However, this extracellular contribution can be tentatively minimized when considering signal attenuation at high diffusion-weighting only, where the extracellular contribution may be expected to have mostly vanished (Pfeuffer et al., 2000). When doing so, lactate diffusion in the control group appears very similar to that of glial/astrocytic metabolites (in particular myo-inositol), with stronger signal attenuation reflecting larger compartment diameter, suggesting lactate is predominantly compartmentalized in astrocytes. This is in line with a recent study using the *Laconic* FRET sensor in the mouse brain (Machler et al., 2016), and consistent with the astrocyte-to-neuron lactate shuttle hypothesis (Pellerin and Magistretti, 1994), which requires a larger concentration of lactate in astrocytes as compared to neurons.

In contrast, lactate diffusion appears to be very similar to that of neuronal metabolites (in particular NAA) in the CNTF group (**Fig. 5B** and **5C**), likely reflecting some major remodeling of lactate cellular compartmentation. Increased oxidative metabolism in reactive astrocytes has been shown to be associated in particular with overexpression of β-hydroxybutyrate dehydrogenase (BDH), providing an alternative source of acetyl-CoA by metabolizing ketone bodies (Escartin et al., 2007) (this might explain the presence of the increased peak at 1.2 ppm on the macromolecule spectrum of the CNTF group, presumably corresponding to more β-hydroxybutyrate interacting with BDH). Moreover, decreased glucose uptake and lactate dehydrogenase activity have also been reported, pointing towards an overall decreased glycolytic activity in CNTF astrocytes. In that context, all pyruvate/lactate produced by astrocytes is likely to be locally consumed to feed increased oxidative metabolism, rather than accumulating in astrocytes; to supplement the deficiency of astrocytic lactate production, neurons may adjust their metabolism and synthesize more lactate, thus resulting in elevated lactate concentrations in neurons.

Although demanding further investigation, these data and their interpretation suggest that DW-MRS may allow non-invasively probing cellular compartmentation of lactate. This would open unprecedented possibilities to study lactate metabolism and astrocyte-neuron interactions in various contexts, such as brain activity (Pellerin and Magistretti, 1994), synaptic plasticity and long-term memory (Magistretti, 2014; Newman et al., 2011; Pellerin and Magistretti, 2012; Suzuki et al., 2011), and Alzheimer’s disease (Demetrius et al., 2014; Harris et al., 2016).

## DISCUSSION

The primary finding of this work is the identification of myo-inositol as the intracellular metabolite whose diffusion is the most sensitive to astrocytic morphological changes. Actually, among intracellular metabolites, significant diffusion alterations could only be detected for myoinositol. Although its role in the brain is still poorly understood, myo-inositol has been proposed as an astrocytic osmolite (Isaacks et al., 1999; Strange et al., 1994; Thurston et al., 1989), and increased myo-inositol concentration is generally associated with astrogliosis. Our results indeed show a strong increase in myo-inositol concentration in the CNTF group. In the light of this work we can, at least to some extent, ascribe this increased myo-inositol content to the increased volume fraction occupied by astrocytes in the volume of interest, due to astrocytic hypertrophy. Because myo-inositol diffusion is altered in the context of hypertrophic astrocytes, it is very tempting to consider myo-inositol compartment shape as a proxy for astrocytic morphology. It turns out that soma radii and process lengths and radii estimated by modeling myo-inositol diffusion are indeed very consistent with actual astrocytic morphology: the agreement between parameters extracted from DW-MRS modeling and confocal microscopy is in general excellent, except for the absolute length of segments. The fact that DW-MRS and confocal microscopy do not yield identical segment length might have different explanations. On one hand, diffusion models are simplified approximations of reality, and as such cannot be anticipated to perfectly reflect cell morphology. In the same vein, the astrocytic specificity of myo-inositol is certainly not perfect, i.e. myo-inositol diffusion might to some extent also reflect the morphology of other cell types where myo-inositol might be present at detectable. On the other hand, confocal microscopy samples only a few cells which might not necessarily represent the whole cell population within the spectroscopic volume of interest. Furthermore it cannot access the extremity of astrocytic processes if they go below optical resolution thus presumably underestimating the length of the thinnest segments. Finally, confocal microscopy is an *ex vivo* technique requiring tissue preparation which usually results in some shrinkage and cutting of histological sections; since in the present study only full astrocytes were analyzed, the longest segments / processes extending beyond slice thickness may have been systematically excluded, resulting in underestimated length. Whatever the origin of this discrepancy, myo-inositol diffusion still appears to be extremely sensitive to astrocyte reactivity, consistently with astrocytic hypertrophy.

The fact that other intracellular metabolites do not exhibit significant diffusion variations, in contrast to myo-inositol, is a strong indication that their astrocytic fraction is small and does not dominate their MRS signal. While this may not be surprising for neuronal metabolites such as NAA and glutamate, or even for non-specific metabolites such as total creatine or taurine, for which it can be conceived that the neuronal contribution dominates, this is rather conflicting with some previous reports of choline compounds being highly abundant in astrocytes cells. A possible explanation might be that astrocytes come in two distinct sub-populations, one being enriched with myo-inositol and becoming hypertrophic, the other being enriched with choline compounds and preserving its morphology.

In conclusion, specific alterations of intracellular metabolite diffusion could be measured and related, for the first time and quantitatively, to cell-specific morphological alteration as independently assessed by microscopy. More specifically, this work establishes DW-MRS as a unique tool to non-invasively monitor astrocytic structural alterations *via* specific variations of myo-inositol diffusion. A complementary step in the future would be to detect specific variations of the diffusion of neuronal metabolites (NAA, Glu) in a context of altered neuronal morphology. Beyond cell structure, this work also suggests that DW-MRS may yield new insights into lactate cellular compartmentation, which is tightly related to the important but still controversial notion of astrocyte-to-neuron lactate shuttling (Diaz-Garcia et al., 2017; Suzuki et al., 2011). Since no fundamental barrier prevents implementing our approach on clinical MRI scanners, the strategy developed here may provide unprecedented access to astrocyte reactivity in the Human brain, especially now that stronger clinical gradients are becoming available (e.g. up to 300 mT/m along each axis for the Human Connectome Project). This could lead to better understanding of the role and status of astrocytes in the healthy and diseased Human brain, and allow monitoring the effect of treatments modulating astrocyte reactivity. Although the present implementation only deals with single-volume detection, recent and promising developments in diffusion-weighted spectroscopic imaging (Bito et al., 2015; Ercan et al., 2015; Fotso et al., 2017) let us foresee the possibility to quantitatively map myo-inositol diffusion, opening the way towards cell-specific microstructure imaging throughout the Human brain.

## MATERIALS AND METHODS

### Mouse model of astrocyte reactivity

All experimental protocols were reviewed and approved by the local ethics committee (CETEA N°44) and submitted to the French Ministry of Education and Research (approval: APAFIS#4554-2016031709173407). They were performed in a facility authorized by local authorities (authorization #B92-032-02), in strict accordance with recommendations of the European Union (2010-63/EEC). All efforts were made to minimize animal suffering and animal care was supervised by veterinarians and animal technicians. Mice were housed under standard environmental conditions (12-hour light-dark cycle, temperature: 22 ± 1°C and humidity: 50%) with ad libitum access to food and water. Self-inactivated lentiviral vectors (LV) pseudotyped with the vesicular stomatitis virus envelope were produced and titrated as described in (Escartin et al., 2006). They encode the human ciliary neurotrophic factor (CNTF) cDNA with the immunoglobulin export sequence or β-galactosidase (LacZ) cDNA under the phosphoglycerate kinase I promoter, and specifically target neurons. To visualize astrocyte morphology with confocal laser microscopy, we used LV encoding green fluorescence protein (GFP) and pseudotyped with the Mokola envelope to target astrocytes. These LV also bear the Mir124 target sequence to repress GFP expression in neurons (Colin et al., 2009). Two-month-old male C57bl6 mice (Charles River, France) were anesthetized with an *i.p* injection of ketamine (100 mg/kg) and xylazine (10 mg/kg). Lidocaine (7 mg/kg) was injected subcutaneously at the incision site, 10 min before injection. LV were diluted in PBS with 1% bovine serum albumin. Three types of striatal injections were performed: 1) bilateral 1 μl-injections of LV-CNTF (83 ng of p24 per site), performed on ten mice forming the “CNTF group” for MRS experiments; 2) bilateral 1 μl-injections of LV-LacZ (83 ng of p24 per site), performed on ten mice forming the “control group” for MRS experiments; and 3) 1.4μl-injections of LV-LacZ + LV-GFP in the left striatum and LV-CNTF + LV-GFP in the right striatum (83 ng of p24 of LV-CNTF or LV-LacZ + 17 ng p24 LV-GFP), performed on five mice for confocal microscopy. Anesthetized mice were placed on the stereotactic frame and were injected with LV dilutions at a rate of 0.2 μl/min with the following coordinates: anteroposterior = +1 mm; lateral +/-2 mm from bregma and ventral = −2.5 mm from the dura, tooth bar set at 0 mm. At the end of injection, the needle was left in place for 5 min before being slowly removed and the skin was sutured. We showed previously that CNTF effects are stable over time(Carrillo-de Sauvage et al., 2015; Escartin et al., 2006).

### MRI and MRS experiments

Mice were scanned 6 weeks after injection on an 11.7 T Bruker horizontal scanner running with Paravision 6.0.1 (Bruker, Ettlingen, Germany), with maximal gradient strength Gmax=752 mT/m on each axis and 100-μs rise time. A quadrature surface cryoprobe (Bruker, Ettlingen, Germany) was used for radiofrequency transmission and reception. Animals were maintained on a stereotaxic bed with a bite and two ear bars. They were anesthetized with 1-1.3% isoflurane in a 1:1 mixture of air and dioxygen (1 L/min total flow rate). Mice temperature was monitored with an endorectal probe and maintained at 37°C with regulated water flow, and respiratory rate was continuously monitored using PC SAM software (Small Animal Instruments, Inc., Stony Brook, NY) during scanning.

A spectroscopic volume of interest (6.5×3×2.8=56 mm^3^) was placed around the striatum (**Fig. 1B**), and signal was acquired with our recent STE-LASER sequence (Ligneul et al., 2017), designed to provide excellent localization while avoiding cross-terms between diffusion gradients and selection gradients. The echo time was TE=33.4 ms, and diffusion gradient duration was set at δ=3 ms. Water suppression was achieved with a VAPOR module and an additional water suppression pulse at the end of the mixing time. The full set of DW-MRS measurements was performed in each mouse. First, spectra at different diffusion-weightings (*b*=0.02, 3.02, 6, 10, 20, 30 and 50 ms/μm^2^, TR=2000 ms, 128 repeats) were acquired at *t*_*d*_=53.2 ms. Then, spectra were acquired at longer *t*_*d*_ for two different *b*-values: *b*_0_ corresponding to the minimal diffusion-weighting achievable while keeping sufficient diffusion gradient strength for efficient spoiling, and *b*=*b*_0_+3 ms/μm^2^ to extract the apparent diffusion coefficient of metabolites according to 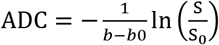. TR was different for the different *t*_*d*_ to allow for similar recovery delay after the LASER localization block. The number of repetitions was increased at the two longest diffusion times to get sufficient SNR. These acquisition parameters for the different *t*_*d*_ are summarized in **Supplementary Table S4**. A macromolecule spectrum was acquired for each mouse (TR=4000 ms, 256 averages). In addition to a double inversion recovery module (TI1=2200 ms, TI2=730 ms) placed prior to the STE-LASER sequence, *b*=10 ms/μm^2^ was applied to reduce residual metabolite peaks. Diffusion MRI was also performed to evaluate its sensitivity to astrocyte reactivity. Images were acquired using 2D spin-echo EPI (TE/TR=30/3200 ms, six segments, 400-kHz acquisition bandwidth), with an 18×18 mm^2^ field-of-view spanned by 160×160 pixels and ten 0.5-mm slices. Diffusion gradient duration was δ=2 ms and gradient separation was Δ=20 ms. In addition to the *b*=0 image, thirty gradient directions were acquired at two *b*-values (1 ms/μm^2^ and 2 ms/μm^2^) to allow for tensor (DTI) and Kurtosis (DKI) analysis. Fractional anisotropy (FA) and mean diffusivity (MD) maps could be reconstructed from DTI and DKI analysis (**Supplementary Fig. S1**). Kurtosis map was also reconstructed from DKI. A region of interest was drawn in the striatum to compute the average value for each of these parameters.

### MRS processing and quantification

Scan-to-scan phase correction was performed on metabolite signal before summing individual scans on Matlab, to correct for incoherent averaging leading to artefactual signal loss. Eddy current correction was achieved using water reference. Spectra were analyzed with LCModel (Provencher, 1993), using a basis set generated for each TM with home-made routines based on the density matrix formalism. Signal could be reliably quantified according to our quality standards (Cramér-Rao lower bounds CRLB<5%) for NAA, tCr, tCho, Glu, Ins, and Tau at all *b*-values and *t*_*d*_. Individual macromolecule spectra acquired in mice were summed with respect to their group and then included in LCModel basis sets associated to the CNTF and control groups. Metabolite concentrations were evaluated from spectra acquired at *t*_*d*_=53.2 ms / *b*=0.020 ms/μm^2^. For this parameter set NAA, tCr, tCho, Glu, Ins, Tau, Gln, GABA and Lac were reliably quantified (CRLB < 5%). As a preliminary check, we summed all spectra of the control group and all spectra of the CNTF group to search for a stable metabolite in both groups to serve as internal reference. Absolute macromolecule signal (peak at 0.9 ppm) appeared to be very stable (see **Fig. 1C**), confirming the stability of the amount of tissue in the volume of interest. Absolute tCr signal was also found to be very stable between both groups (**Fig. 1B**), therefore we used it as an internal reference standard (fixed at 8 mM in both groups) for quantification of metabolite concentration in all individual spectra.

### Statistical analysis of MRS data

For metabolite levels measured in MRS experiments, statistical significance of the effect of the group (CNTF or control) on metabolites concentrations (only those quantified with Cramér-Rao lower bounds <5%) was assessed by an unpaired Student’s t-test. To compensate for the type I error due to multiple comparisons, p-values were adjusted by a Bonferroni correction over the eight metabolites considered. In DW-MRS experiments, we wanted to evaluate the statistical significance of the effect of the group (CNTF or control) on the functional form of the diffusion, i.e. signal attenuation over *b* or on ADC over *t*_*d*_, as this functional form carries the information about the underlying microstructure. This was assessed by an analysis of variance (ANOVA) with one between-subject factor (group CNTF or control) and one within-subject factor (*b*-value or *t*_*d*_) (**Supplementary Table S5**). Data homoscedasticity was confirmed by a Barlett test. After identifying metabolites exhibiting a significant difference in diffusion properties between groups, i.e. a group\**b* or group\**t*_*d*_ interaction (which was the case only for myo-inositol), we ran unpaired Student’s t-test as *post hoc* tests on each value of the within-subject factor, i.e. on each point of myo-inositol diffusion curves (signal attenuation as a function of *b* or ADC as a function of *t*_*d*_). To compensate for the type I error due to multiple comparisons over the six levels of each factor (six non-zero *b*-values, and six *t*_*d*_), p-values were adjusted by a Bonferroni correction with *n*=6.

### Diffusion modeling for analysis of high-*b* data to evaluate the radius of fibers and soma

To analyze the DW-MRS data and estimate the size of cellular soma and projections, we used our recently introduced cylinder model (Palombo et al., 2017b), with a small correction to account for soma contribution. Specifically, cellular projections can be described in first approximation as a collection of long cylinders (the “fibers”), with radius *R*_*fiber*_, and intracellular diffusivity *D*_*intra*_. We assume the cylinders are randomly oriented to calculate the signal attenuation *S*_*fiber*_ in the narrow pulse approximation, as described in (Palombo et al., 2017b). Here we consider the cellular volume fraction occupied by astrocytic processes to be *f*, the remaining intracellular fraction 1-*f* being occupied by the soma. Studies carried on different species and with different methods (Chvatal et al., 2007) report fibers representing 80-85% of the total astrocytic volume. Our own confocal measurements (see details below) yield *f*~77% in control and ~71% in CNTF. In line with these estimates, in our model we set *f*=80%. The DW-MRS signal originating from cellular soma, *S*_*soma*_, is modeled as the signal arising from molecular diffusion restricted within a sphere (Linse and Soderman, 1995) of radius *R*_*soma*_ and intracellular diffusivity *D*_*intra*_, and it contributes to the total DW-MRS signal *S* with a weight 1-*f* = 20%, in our model. In the end:

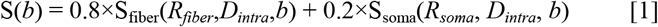

Equation 1 was fitted to myo-inositol signal attenuation as measured *in vivo* and averaged over all animals from each group, to estimate *R*_*soma*_ and *R*_*fiber*_ (with *D*_*intra*_ let as a free parameter as well).

Fitting was performed by using a nonlinear least-square regression, based on the trust-region-reflective algorithm implemented in MATLAB (The MathWorks, Natick, Massachusetts, USA). The error on estimated parameters was evaluated using a Monte Carlo (MC) approach (*n*=2,500 draws). For fig. each draw, random noise (whose standard deviation s.d. was estimated from the difference between the best fit and the experimental data) was generated and added to the best fit to generate a new data set, which could be analyzed using the model. Statistical difference between control and CNTF values was assessed by permutation analysis of the parameter distribution estimated from the 2,500 MC draws using unpaired two-tailed t-test (with Bonferroni correction for the the two structural parameters of interest *R*_*fiber*_ and *R*_*soma*_). The results of this fitting procedure are reported in **Supplementary Fig. S8** and **Supplementary Table S2**.

### Diffusion modeling and machine learning for analysis of long-*t*_*d*_ data and evaluation of long-range cellular morphology

The ADC time-dependency of brain metabolites was analyzed using the computational framework introduced in (Palombo et al., 2016). However, unlike in (Palombo et al., 2016), a dictionary-based approach was used here for better computational efficiency. An efficient code was implemented in Matlab to parallelize this huge amount of computation on a GPU device (NVIDIA Tesla K20c). The whole procedure required about 3 minutes for each set of morphometric parameters, and about 400 hours to build the full database by varying D_intra_, mean N_branch_ and mean L_segment_. The number of processes radiating from the soma has no effect on ADC time-dependency (Palombo et al., 2016), and was therefore not varied (fixed to 5±2). To limit the degrees of freedom, we also fixed the standard deviations on the successive number of embranchments and on segment length (SDN_branch_=2 and SDL_segment_=10 μm), since the effect on ADC(*t*_*d*_) is much smaller than the effect of mean N_branch_ and L_segment_.

To learn the mapping between the substrate parameters and ADC time-dependency, a random forest regression was used (Nedjati-Gilani et al., 2017). We used a widely used and freely available random forest regressor in the sci-kit learn python toolkit (Pedregosa et al., 2011) with 100 trees and a maximum depth of 20 (preliminary experiments found that we obtain diminishing returns in accuracy above these values). More in-depth details of the implementation are available on the scikit-learn website (http://scikit-learn.org/). The forest was then trained on 6,400 simulated datasets, with the remaining 1,400 previously unseen feature vectors used for testing. Following the training and testing stages, we used the forest regressor on the ADC time-dependency as measured *in vivo* (averaged over all animals of each group) (**Supplementary Fig. S9**). Stability of the estimated parameters with respect to experimental noise was evaluated using a Monte Carlo approach (2,500 draws of noised datasets). At each Monte Carlo iteration, random Gaussian noise was generated with s.d. comparable to experimental s.d. on ADC (~15%) and added to the reference dataset to generate a new dataset, which was analyzed using the same pipeline. For each of the three parameters, a distribution of 2,500 estimated values was obtained, and the best estimates and its uncertainty were then computed as the mean and s.d. of this distribution. These values are reported in **Supplementary Table S3**, where statistical difference between control and CNTF values was assessed by permutation analysis of the parameter distribution estimated from the 2,500 MC draws using unpaired two-tailed t-test (with Bonferroni correction for the the two structural parameters of interest Branch Order and L_segment_).

### Histology and confocal microscopy acquisitions

Transcardial perfusion fixation of anesthetized mice, using cold 4% paraformaldehyde, was performed to get brain samples. Collected brains were kept at 4°C in paraformaldehyde solution at least one night and then transferred to PBS. 100 - 120 μm cortical sections of mouse brains were obtained with a vibrating-blade microtome (Leica) and then mounted in Vectashield mounting medium (Vector Labs). Isolated or small groups of astrocytes were imagined using a Leica SP8 confocal system equipped with a broadband white-light laser, with an oil immersion objective 63X, 1.4 NA. The excitation laser line was setup to 489 nm for eGFP excitation at 3% of intensity. Sequential scan mode imaging was used setting up the HyD photodetector to 498-570 nm for eGFP labeling. For all images a 1 airy unit pinhole was setup based on a 510 wavelength emission and the XY resolution and Z-step size was automatically optimized based on the Nyquist theorem. In all cases the white laser power was set to 70%, and the acquisition speed of the laser scan was 600 Hz.

### Confocal microscopy image analysis and astrocytes reconstruction

Image analysis was performed using Fiji (Schindelin et al., 2012) Vaa3D (Peng et al., 2014), Matlab (MathWorks, USA), and Autoquant X3.1 (Media Cybernetics, USA). To reduce the workload due to the large size of original images (≈4 GB), regions of interest were cropped and each ROI was slice aligned in the X and Y axis and deconvoluted using Autoquant X3.1; through the blind adaptive PSF method with ten Iterations and low noise level. Then, images were adjusted in contrast using Fiji. Once in Vaa3D, the reconstruction of the astrocytes was made using the Vaa3D-Neuron2 Auto Tracing Based on APP2 plugin; basically, one marker was placed on the soma center of the single astrocyte and the plugin was run setting an auto-thresholding for the background, activating the option to get the radius from 2D and adjusting the length threshold until get a proper reconstruction, all the other options were settled as default (see **Supplementary Fig. S2** to **S5**). Then, the structure was sorted using the “sort neuron” SWC plugin. To label the soma section, a six-face ROI was created around the user defined volume and the segments inside that volume were labeled as soma, then each file was reviewed to correct wrong and missed segments. As Vaa3D only analyzes the SWC files in voxel units, the SWC were exported to Matlab using the TREES toolbox (Cuntz et al., 2010) and then the model was resized to the real dimensions from the initial acquisition. Finally, information from the model was extracted using L-measure (Scorcioni et al., 2008). The approach used to estimate the radius of the confocal reconstructions was to obtain the approximate circle of cross section area labeled in the XY plane based on the image for each marker of the reconstruction.

### Comparison between real and synthetic astrocytes: cellular branching structure representation and group statistics

When astrocytic fibers are regarded as graphs, their branching structure can be well described with the corresponding directed adjacency matrix, a quadratic matrix of size N×N where N is the number of nodes. However, the adjacency matrix formalism for cell-graphs does not represent one and the same graph in a unique way. The same is true for the.swc format used to reconstruct the cell morphology from confocal microscopy. A more constrained representation is given by the BCT formalism (Eeckman et al., 1994). The nodes are labeled sequentially so that the nodes of a subregion of the graph remain in sequence, and within each subregion of the graph parent nodes always precede their respective daughter nodes. Nodes with no daughter node are termed terminal, “T”; nodes with only one daugther node are termed continuation, “C”; and nodes with more than one daughter node are termed branch, “B”. Both real and synthetic cell morphologies are represented in the BCT formalism using the TREES toolbox (Cuntz et al., 2010). However, before comparing real structures with synthetic ones, some simplifications have to be done in order to compare structure using similar approximations. In the modeling strategy introduced in the previous section, a couple of approximations have been done: (i) small terminal segments radiating from a parent branch (leaflets-like structures, i.e. terminal segments less than 5 μm in length) are not considered relevant on the ADC time-dependency (Palombo et al., 2017a); (ii) there are no C segments, which means undulations and curvature of the cell branch is not modeled and the branch joining two consecutive nodes is just a straight segment, representing the euclidean distance between the two nodes in 3D. According to these two approximations, astrocytes reconstructed from confocal microscopy were simplified, preserving the original topological characteristics, as shown in **Supplementary Fig. S10**. Moreover, in order to compare real and synthetic cell structures, we used the Branch Order rather than the number of consecutive bifurcation, N_branch_, because the first is used to describe the cell-graph structure in computational neuroscience (Cuntz et al., 2010), while Nbranch is an *ad hoc* parameter that we introduced to control the complexity of our cell-graphs. Note that we empirically found a simple relationship between Branch Order and N_branch_: Branch Order ~ N_branch_ + 2.

Two groups of 20 cell structures reconstructed from confocal microscopy from control and CNTF mice, named “real control” and “real CNTF” respectively, were compared with two groups of similar size from synthetic cell structures as estimated from DW-MRS in control and CNTF mice, named “synth control” and “synth CNTF” respectively. Statistics for L_segment_ and Branch Order in all groups are reported in **Supplementary Table S3**.

## Acknowledgements

This research was supported by the European Research Council (grant agreement 336331 - INCELL project, awarded to J.V.). M.P. acknowledges support from EPSRC grant EP/N018702/1. The 11.7 T MRI scanner and the confocal microscope were funded by a grant from “Investissements d’Avenir - ANR-11-INBS-0011 - NeurATRIS: A Translational Research Infrastructure for Biotherapies in Neurosciences”.

## Supplementary table legends

**Table S1**. Metabolite concentrations in control and CNTF groups (mean±s.d, in mM), using tCr as an internal reference at 8 mM.

**Table S2**. Fiber and soma radii estimated from confocal microscopy images and from DW-MRS signal attenuation as a function of *b* for myo-inositol, in control and CNTF mice. The values from *ex vivo* confocal microscopy (“Real”) represent the average values over the population of 20 cell structures reconstructed from confocal microscopy images (s.d. in brackets). The values from *in vivo* DW-MRS (“Synthetic”) represent the estimated values by fitting equation 1 to the experimental myo-inositol data, with s.d. being evaluated using a Monte Carlo (MC) approach (*n*=2,500 draws). Statistical difference between control and CNTF values was assessed by permutation analysis of the parameter distribution estimated from the 2,500 MC draws using unpaired two-tailed t-test with Bonferroni correction (** =*p*<0.01; *** =*p*<0.001).

**Table S3**. Mean and s.d. of the long-range morphometric statistics (L_segment_ and Branch Order) estimated on 20 cells, reconstructed either from confocal microscopy images (“real”) or when fitting the ADC time-dependency of myo-inositol with the computational framework described in the text (“synthetic”). Statistical difference between control and CNTF values was assessed by permutation analysis of the parameter distribution estimated from the 2,500 MC draws using unpaired two-tailed t-test with Bonferroni correction (*** =*p*<0.001).

**Table S4**. DW-MRS acquisition parameters for long-*t*_*d*_ measurements.

**Table S5**. ANOVA group\**b* and group\**t*_*d*_ interactions for the different intracellular metabolites quantified with CRLB<5% at all *b*/*t*_*d*_.

### Supplementary figure legends

**Figure S1**. Masked parameter maps as derived from diffusion MRI, exemplified here in one mouse. Parameters maps derived from Kurtosis analysis are shown in upper row: fractional anisotropy (DKI - FA), mean diffusivity (DKI - MD, in μm^2^/ms) and Kurtosis (DKI - Kurtosis). Parameters maps derived from tensor analysis are shown in lower row: fractional anisotropy (DTI - FA) and mean diffusivity (DTI - MD, in μm^2^/ms). In the lower right corner, the average values as measured in the striatum for these five parameters, in control and CNTF mice (*n*=10 per group, error bars stand for s.d.).

**Figure S2**. Examples of real astrocytes from control mice. (**A-J**) Confocal images maximum projection of different GFP-labeled striatal astrocytes of mice in which neurons secreted β-Galactosidase as control; in all cases scale bar correspond to 10 μm. (**A’-J’**) Reconstructions based on the confocal images of each selected astrocytes.

**Figure S3**. Other examples of real astrocytes from control mice. (**A-J**) Confocal images maximum projection of different GFP-labeled striatal astrocytes of mice in which neurons secreted β-galactosidase as control; in all cases scale bar correspond to 10 μm. (**A’-J’**) Reconstructions based on the confocal images of each selected astrocytes.

**Figure S4**. Examples of real astrocytes from CNTF mice. (**A-J**) Confocal images maximum projection of different GFP-labeled striatal astrocytes of mice in which neurons secreted CNTF; in all cases scale bar correspond to 10 μm. (**A’-J’**) Reconstructions based on the confocal images of each selected astrocyte.

**Figure S5**. Other examples of real astrocytes from CNTF mice. (**A-J**) Confocal images maximum projection of different GFP-labeled striatal astrocytes of mice in which neurons secreted CNTF; in all cases scale bar correspond to 10 μm. (**A’-J’**) Reconstructions based on the confocal images of each selected astrocyte.

**Figure S6**. Probability density distributions (pdf) and corresponding box-and-whisker plots of the long-range morphometric statistics (L_segment_ and Branch Order) estimated in control and CNTF groups from 20 cells, either reconstructed from confocal microscopy images (“real”) or from modeling the ADC time-dependency of myo-inositol (“synth”).

**Figure S7**. Stackplot of spectra, zoomed around the lactate peak, as a function of *b* (at fixed *t*_*d*_=53.2 ms). Here spectra at each *b* are averaged over the 10 mice per group to increase signal-to-noise ratio and thus strikingly illustrate the different diffusion behavior in the two groups: while lactate almost disappears at high *b* in the control group, it remains relatively high in the CNTF group. The macromolecule spectra of each group (in color) are also superimposed on the spectra at the highest *b*, to show that the elevated lactate peak in CNTF is not an artefact due to the elevated β-hydroxybutyrate at 1.2 ppm.

**Figure S8**. Fit of the high-*b* signal attenuation for myo-inositol in control (blue) and CNTF (orange) mice, to estimate the radius of astrocytic soma and fibers, using Equation 1.

**Figure S9**. Fit of the long-*t*_*d*_ ADC for myo-inositol data in control (blue) and CNTF (orange) mice, to estimate fiber branching and length using machine learning based on numerical simulations.

**Figure S10**. Schematic description of the steps performed to simplify astrocytic structures reconstructed from confocal microscopy and make them comparable with branch structures derived from DW-MRS modeling. (**A**) Example of one astrocyte reconstructed from confocal microscopy resolution. (**B**) As our long-*t*_*d*_ DW-MRS does not account for leaflet-like structures, such structures are deleted from the reconstruction. (**C-D**) Fiber undulations are not taken into account in our models, and hence are also “removed”: for that, nodes of continuity (black points) are identified (**C**) and joined together (**D**). This step converts the “curved” cell structure into a “straightened” structure that presents similar topological features (like the number and spatial distribution of nodes, nodes connectivity, spatial extension of the graph).

